# Presenilin-dependent regulation of tau pathology via the autophagy/proteasome pathway

**DOI:** 10.1101/2023.12.21.572822

**Authors:** Anna del Ser-Badia, Carlos M. Soto-Faguás, Rebeca Vecino, José Rodríguez-Alvarez, Carlos Vicario, Carlos A. Saura

**Affiliations:** Institut de Neurociències, Department de Bioquímica i Biologia Molecular, Universitat Autònoma de Barcelona, Barcelona 08193, Spain; Centro de Investigación Biomédica en Red Enfermedades Neurodegenerativas (CIBERNED), Instituto de Salud Carlos III (ISCIII), Spain; Instituto Cajal, Consejo Superior de Investigaciones Científicas (CSIC), Madrid 28002, Spain

**Keywords:** Alzheimer’s disease, γ-secretase, neurodegeneration, proteostasis, tauopathies

## Abstract

Autosomal dominant inherited mutations in the presenilin (*PS*/*PSEN*) genes cause early-onset familial Alzheimer’s disease (AD) by enhancing cerebral accumulation of amyloid-β (Aβ) and microtubule-associated protein tau, although the precise cellular mechanisms by which PS dysfunction drives neuronal tau pathology remain still unclear. Here, we investigated the mechanisms linking PS/γ-secretase-dependent tau pathology and autophagy by using molecular, imaging and pathological approaches in brains, fibroblasts and induced pluripotent stem cells (iPSCs)-derived neurons from mutant *PSEN1* carriers, as well as in a novel tauopathy mouse model lacking *PS* in glutamatergic neurons. We found colocalization of phosphorylated tau with the autophagy marker p62 in the hippocampus of tauopathy patients with *PSEN1* mutations, corticobasal degeneration and Pick’s disease. Remarkably, disrupted autophagic clearance of pathological tau was evidenced by increased autophagy markers and accumulation of total and AD-associated phosphorylated tau species (pTau 181, 202, 217) in hippocampal lysates and autophagosomes of familial AD-linked *PSEN1* patients and *PS*-deficient tau transgenic mice. Human iPSC-derived neurons harboring the familial AD-linked *PSEN1* G206D mutation are less sensitive to autophagy inhibition, reduce tau release and accumulate intracellular tau oligomers. Human primary fibroblasts from *PSEN1* G206D and/or L286P carriers show elevated LC3 and autolysosomes indicating that these familial AD-linked *PSEN1* mutations disrupt autophagy flux. PS is required for efficient autophagy-mediated tau degradation in neurons through a dual mechanism involving autophagy induction via blockage of Akt/PRAS40-dependent mTORC1 activation and promoting autophagosome/lysosome fusion. Surprisingly, pharmacological proteasome inhibition decreases tau accumulation in neurons by promoting tau release through a mechanism that requires functional *PS.* In conclusion, PS is required for autophagy/proteasome-mediated tau elimination in neurons, while familial AD-linked *PSEN* mutations cause progressive tau pathology by disrupting autophagy. These findings may impact on the development of new therapeutic targets for tauopathy dementias.

## Background

Alzheimer’s disease (AD) is the most prevalent memory disorder in the elderly affecting 57 million people worldwide. AD is characterized by cerebral accumulation of amyloid plaques, containing amyloid-β (Aβ) peptides, and neurofibrillary tangles (NFTs) composed of aggregated hyperphosphorylated microtubule-associated protein tau (MAPT) (Ittner and Gotz, 2011; Spires-Jones and Hyman, 2014). Changes in cerebral Aβ, tau and glucose metabolism are detected decades before disease clinical onset (McDade et al., 2018), and elevated phosphorylated tau (pTau 181, 217 and 231) in CSF and plasma are associated with cognitive decline and conversion to dementia being potentially useful biomarkers for clinical diagnosis (Janelidze et al., 2020; Simrén et al., 2021; Ashton et al., 2022; Frank et al., 2022). Indeed, cerebral tau pathology correlates tightly with cognitive decline appearing in the temporal lobe and then progresses to limbic and association cortices (Arriagada et al., 1992; Gomez-Isla et al., 1997). Although Aβ and tau act synergistically in AD (Roberson et al., 2007; Ittner et al., 2010; Pickett et al., 2019), tau also causes neurodegeneration independently of Aβ (DeVos et al., 2018), as evidenced by tau pathology in frontotemporal dementias (FTD), including corticobasal degeneration (CBD), Pick’s disease (PiD) and progressive supranuclear palsy (PSP) (Goedert et al., 2017).

Autosomal dominant-inherited mutations in the *presenilin* (*PS/PSEN*) genes, encoding the catalytic components of γ-secretase presenilin-1 (*PSEN1*) and presenilin-2 (*PSEN2*), account for the majority of familial AD (FAD) cases (De Strooper et al., 2012). These pathogenic mutations increase cerebral levels of toxic Aβ peptides, which determine the pathogenicity and predict disease onset (Petit et al., 2022), and tau-associated neurites causing accelerated disease progression (Gomez-Isla et al., 1999; Sudo et al., 2005; Woodhouse et al., 2009). Symptomatic FAD-linked *PSEN1* carriers show changes in CSF biomarkers, including lower Aβ42 and elevated levels of tau, pTau 181 and neurofilament (Sanchez-Valle et al., 2018). FAD-linked *PSEN* mutations also increase tau phosphorylation, amyloid deposition, synaptic dysfunction and neurodegeneration in mice (Dewachter et al., 2008; Xia et al., 2015; Ochalek et al., 2017). These mutations interfere with γ/ε-secretase-dependent and -independent PS biological functions revealing a possible loss of function mechanism due to FAD-linked *PSEN* mutations (Lleó and Saura, 2011). Consistently, PS deficiency in neurons results in age-dependent neurodegeneration accompanied by tau phosphorylation in *PS* conditional knockout (cKO) mice (Saura et al., 2004; Watanabe et al., 2014; Soto-Faguas et al., 2021). Remarkably, *PSEN1* mutations enhance tau accumulation in the absence of Aβ pathology in rare cases of familial FTD, atypical dementia with parkinsonism and dementia with Lewy bodies (Raux et al., 2000; Dermaut et al., 2004b; Bernardi et al., 2009), which suggests that pathological tau can accumulate independently of Aβ (Hutton, 2004; Larner and Doran, 2006). Although loss of PS/γ-secretase recapitulates key features of tauopathies, **the PS-dependent neuronal mechanisms leading to tau pathology in neurodegeneration are still unknown**.

Macroautophagy, a catabolic cellular process in which proteins and organelles are targeted to autophagosomes for degradation, plays a central role in neurodegenerative diseases, including tauopathies (Menzies et al., 2017). The accumulation of autophagic vacuoles and autolysosomes containing tau in sporadic and amyloid precursor protein (APP)-linked familial AD, CBD and PSP cases suggests disrupted autophagy-lysosomal degradation in tauopathies (Nixon et al., 2005; Boland et al., 2008; Piras et al., 2016). Tau is eliminated via the autophagy/lysosomal system, whereas pathological tau disrupts macroautophagy and chaperone-mediated autophagy (CMA) contributing to toxicity in AD (Kruger et al., 2012; Jo et al., 2014; Caballero et al., 2018; Caballero et al., 2021). Indeed, autophagy activators potentiate tau clearance and mitigate tau toxicity (Kruger et al., 2012; Schaeffer et al., 2012; Caccamo et al., 2013; Ozcelik et al., 2013; Silva et al., 2020; Bourdenx et al., 2021). Synaptic activity also decreases pathological tau by promoting its degradation via the autophagy/lysosomal pathway and increasing its extracellular release (Pooler et al., 2013; Yamada et al., 2014; Akwa et al., 2018). However, **the mechanisms linking autophagy and tau accumulation in tauopathies are unclear**.

PS1 promotes autophagy-mediated proteolysis independently of γ-secretase by facilitating autophagosome-lysosome fusion (Lee et al., 2010; Neely et al., 2011; Lee et al., 2015; Bustos et al., 2017). In agreement, loss of PS function and pathogenic FAD-linked PS mutations impair autophagy and lysosome fusion by deregulating autophagy-related genes and lysosome acidification (Cataldo et al., 2004; Lee et al., 2010; Neely et al., 2011; Dobrowolski et al., 2012; Reddy et al., 2016; Martín-Maestro et al., 2017; Chong et al., 2018; Hung and Livesey, 2018; Fedeli et al., 2019). Besides the critical role of PS in autophagy (see (Oikawa and Walter, 2019) for a recent review), whether PS regulates autophagy-mediated tau clearance in neurodegeneration is still unclear. Here, we investigated the mechanisms of PS-dependent tau pathology by examining the link between tau and autophagy markers in human brains, fibroblasts and iPSC-derived neurons from FAD-linked *PSEN1* carriers, as well as in novel *in vivo* and *in vitro* mouse models of tauopathy lacking PS function. We found that PS regulate proteasome- and autophagy-mediated tau elimination, and that familial AD-linked *PSEN* mutations may drive tau pathology through a loss of function mechanism in autophagy.

## Methods

### Human brain samples

Human post-mortem hippocampus from controls (n=12; no neurodegenerative pathology), FAD-linked *PSEN1* (n=6) and *PSEN2* (n=1) carriers, CBD (n=8) and PiD (n=7) subjects were obtained from the Hospital Clinic-IDIBAPS Biobank (Universidad de Barcelona, Spain). Samples were matched as closely as possible for sex, age and postmortem interval (Table 1). Neuropathology was classified according to Braak staging for neurofibrillary tangles and neuritic plaques (Braak et al., 2006). Experimental procedures were conducted according to the Animal and Human Ethical Committee of the Universitat Autònoma de Barcelona (CEEAH 2896) and Generalitat de Catalunya (DMAH 8787) following the experimental European Union guidelines and regulations (2010/63 and 2016/679).

**Table 1.** Human hippocampal samples used in this study. Samples are classified according to the clinical and neuropathological diagnosis. FAD: Familial Alzheimer’s disease; CBD: corticobasal degeneration; PiD: Pick’s disease; F: female; M: male; PMD: postmortem delay. BC: biochemistry; IHC: immunohistochemistry

| Sample ID | Gender | Age | Clinical Diagnosis | PMD (h:min) | BC | IHC |
| --- | --- | --- | --- | --- | --- | --- |
| A08-42 | F | 64 | Control | 5:00 | ✓ |  |
| A06-189 | F | 46 | Control | 9:35 | ✓ |  |
| A08-54 | F | 55 | Control | 8:30 | ✓ |  |
| A07-128 | M | 43 | Control | 4:35 | ✓ |  |
| A07-5 | M | 56 | Control | 5:00 | ✓ |  |
| A10-17 | M | 52 | Control | 3:00 | ✓ |  |
| A09-145 | M | 44 | Control | 6:40 | ✓ |  |
| 1818 | M | 78 | Control | 5:00 |  | ✓ |
| 1937 | F | 83 | Control | 7:33 |  | ✓ |
| 1949 | M | 86 | Control | 7:25 |  | ✓ |
| 1870 | F | 97 | Control | 7:20 |  | ✓ |
| 2055 | M | 94 | Control | 15:46 |  | ✓ |
| 929 | F | 56 | FAD *PSEN1* P246L | 6:00 | ✓ | ✓ |
| 1241 | F | 48 | FAD *PSEN1* M139T | 16:41 | ✓ | ✓ |
| 1846 | F | 61 | FAD *PSEN2* R62H | 7:00 | ✓ | ✓ |
| 2082 | F | 53 | FAD *PSEN1* L286P | 7:15 | ✓ | ✓ |
| 898 | M | 57 | FAD *PSEN1* M139T | 15:25 | ✓ | ✓ |
| 935 | M | 44 | FAD *PSEN1* E120G | 5:30 | ✓ | ✓ |
| 1134 | M | 53 | FAD *PSEN1* M139T | 5:25 | ✓ | ✓ |
| 1058 | F | 69 | CBD | 12:10 | ✓ | ✓ |
| 1291 | F | 79 | CBD | 10:30 | ✓ | ✓ |
| 1359 | F | 67 | CBD | 6:45 | ✓ | ✓ |
| 1798 | F | 78 | CBD | 5:30 | ✓ | ✓ |
| 1204 | M | 74 | CBD | 14:30 | ✓ | ✓ |
| 1356 | M | 72 | CBD | 7:00 | ✓ | ✓ |
| 1518 | M | 66 | CBD | - | ✓ | ✓ |
| 1751 | M | 59 | CBD | 7:00 | ✓ |  |
| 1248 | F | 41 | PiD | 11:10 | ✓ |  |
| 1493 | F | 75 | PiD | 13:00 | ✓ | ✓ |
| 1681 | F | 69 | PiD | 3:00 | ✓ | ✓ |
| 1352 | M | 74 | PiD | 13:00 | ✓ | ✓ |
| 1719 | M | 73 | PiD | 9:15 | ✓ | ✓ |
| 1789 | M | 78 | PiD | 11:00 | ✓ | ✓ |
| 2064 | F | 70 | PiD | 5:45 |  | ✓ |

### Transgenic mice

*PS* cKO mice (C57BL/6/129 background) lacking *PS2* and *PS1* specifically in forebrain glutamatergic neurons were previously described (Saura et al., 2004). Littermate control (*PS1* f/f; *PS2*+/+ or *PS1* f/f; *PS2*+/−; f:floxed), *PS1* cKO (*PS1* f/f; CaMKIIα-Cre) and *PS* cKO (*PS1* f/f; *PS2*-/-; CaMKIIα-Cre) mice were obtained by crossing floxed *PS1*/*PS2-/-* (*PS1* f/f; *PS2*-/-) or *PS2*+/− (*PS1* f/f; *PS2*+/−) males to heterozygous *PS1* cKO; *PS2*+/− females (*PS1* f/f; *PS2*+/−; CaMKIIα-Cre). We generated novel non-transgenic control (*PS1* f/f;Tau-), tau (*PS1* f/f;Tau+), *PS1* cKO;Tau (*PS1* f/f;CaMKIIα-Cre;Tau+) and *PS* cKO;Tau (*PS1* f/f;*PS2*-/-; CaMKIIα-Cre;Tau+) mice by crossing *PS1* cKO or *PS* cKO with Tau P301S tg (PS19) mice (JAX #008169; C57BL/6). Tau P301S mice express the FTDP-17-linked P301S tau under the neuron-specific prion protein promoter leading to behavior alterations and pTau and neurofibrillary tangles pathologies at 5-8 months (Yoshiyama et al., 2007).

### Primary neuronal and fibroblast cultures

Cortical neurons from embryonic mouse brains (E15.5) were cultured in poly-D-lysine coated dishes containing neurobasal medium supplemented with B27 and glutamine (Life Technologies). Neurons (4 DIV) were transduced with lentivirus (1-2 transducing units/cell) and then lysed (12 DIV) with cold RIPA-DOC buffer (50 mM Tris HCl, pH 7.4, 150 mM sodium chloride, 0.1% SDS, 1% NP-40, 0.5% sodium deoxycholate, 2.5 mM EDTA, 1 mM Na_3_VO_4_, 25 mM sodium fluoride). Lentiviral vectors containing Cre-recombinase and ΔCre-recombinase (control) were previously reported (Watanabe et al., 2009). Primary human skin fibroblasts were plated at 4000-5000 cells/cm^2^ and maintained in complete Dulbecco’s Modified Eagle’s Medium (DMEM). Fibroblasts plated in 24-well dishes were transfected with Lipofectamine 2000 reagent following the manufacturer’s protocol (Thermo Fisher Scientific). Cells were fixed with 4% PFA and analyzed with a Zeiss Axio Examiner LSM700 laser scanning microscope 24 h after transfection. For pharmacological treatments, cultured cortical neurons, primary fibroblasts or iPSCs-derived neurons were kept in half of the medium and incubated with vehicle, chloroquine (10 or 25 μM, 24 h; Sigma-Aldrich) and/or MG132 (1 μM, 24 h; Tocris).

### Differentiation of human iPSCs into hippocampal neurons

Human iPSCs were obtained by reprogramming dermal fibroblasts isolated from skin biopsies of FAD patient carrying the G206D mutation in *PSEN1* gene and healthy subjects (Diaz-Guerra et al., 2019; Rodriguez-Traver et al., 2020). The human iPSCs were grown on vitronectin and treated with small molecules LDN 193189 and A83-01 (Miltenyi Biotec) to induce a neural cell fate as well as with cyclopamine (Stem cell) to inhibit the sonic hedgehog signalling pathway and promote a telencephalic dorsal phenotype. Neural progenitor cells (NPCs) were expanded by adding FGF-2 (Peprotech) and then were seeded at 300,000 cells/cm^2^ in polyornithine (Sigma) and laminin-treated plates (ThermoFisher). DAPT (Tocris), Wnt3a, BDNF, NT-3 (Peprotech) and cAMP (Sigma) were added to promote the generation and differentiation of hippocampal neurons, which were cultured for 60 DIV.

### Quantification of human Aβ and tau

A multiplex assay kit (HNABTMAG-68K, Merck Millipore) was used to quantify Aβ40, Aβ42 and total tau levels in the medium of Tau transgenic neurons and iPSCs-derived neurons. Culture medium was harvested, centrifuged and incubated (O/N, RT) with the detection antibodies and the antibody-immobilized magnetic beads. Then, samples were incubated with streptavidin-phycoerythrin (30 min, RT) and before measurements on a Luminex MAGPIX® System (Merck-Millipore).

### Purification of autophagic fractions from brain

Purification of autophagic fractions from brain samples was performed using a self-established modified protocol originally described for liver (Bains and Singh, 2021). Briefly, human hippocampal tissue (0.1 g) was homogenized in ice-cold 0.25 M sucrose with a Teflon-glass homogenizer and centrifuged 5 min at 2,000 x *g*. The supernatant was centrifuged 12 min at 17,000 x *g* and the resulting pellet, which contains the autophagic fractions, was resuspended in 0.25 M sucrose and 51% Nycodenz (ProteoGenix). A 26%, 24%, 20% and 15% discontinuous Nycodenz gradient was loaded on top of the sample layer, and centrifuged 3 h at 104,300 x *g*. The autophagosome, autolysosome and lysosome fractions were collected from the 15%-20%, 20%-24% and 24%-26% Nycodenz interphases, respectively. Each fraction was diluted with ice-cold 0.25 M sucrose and centrifuged 1 h at 30,000 x *g*. The pellets were resuspended in RIPA-DOC buffer.

### Biochemical analysis

For biochemical analysis, half hippocampi were lysed in cold-lysis buffer (62.5 mM Tris hydrochloride, pH 6.8, 10 % glycerol, 5 % β-mercaptoethanol, 2.3 % sodium dodecyl sulfate [SDS], 5 mM NaF, 100 µM Na_3_VO_4_, 1 mM EDTA, 1mM ethylene glycol tetraacetic acid) containing protease and phosphatase inhibitors and boiled at 100°C (Planet et al., 2001). Proteins were quantified with the Coomassie (Bradford) assay kit (Thermo Fisher Scientific), resolved on SDS-polyacrylamide gel electrophoresis and detected by Western blotting with the following antibodies: rabbit anti-tau (D1M9X; 1:1000, Cell Signaling), mouse anti-tau (TG5; 1:500), mouse anti-phosphorylated tau Thr181 (AT270; 1:200, Thermo Fisher Scientific), Ser202 (CP13; 1:250) or Ser396/404 (PHF-1; 1:250), rabbit anti-phosphorylated tau Thr217 (1:1000, Thermo Fisher Scientific), rabbit anti-LC3B (1:1000, Abcam), mouse anti-p62/SQSTM1 (1:1000, Abcam), rabbit anti-LAMP2A (1:1000, Abcam), rabbit anti-PS1 NTF (1:10000, Calbiochem), mouse anti-β-tubulin (1:20000, Sigma), mouse anti-GAPDH (1:100000; Ambion), mouse anti-GSK3β (1:2500; BD Biosciences), rabbit anti-phosphorylated GSK3β (Ser9, 1:1000; Cell Signaling), goat anti-Akt (1:1000; Santa Cruz Biotechnology), and rabbit anti-phosphorylated Akt (Thr308 and Ser473), total and phosphorylated (Thr172) AMPK, total and phosphorylated (Ser2448) mTOR, phosphorylated PRAS40 (Thr246), total and phosphorylated (Thr389) p70 S6K, total and phosphorylated (Ser235/236) S6 ribosomal protein and pULK1 (Ser757) (all 1:1000, Cell Signaling). Bands detected with secondary antibodies coupled to peroxidase and enhanced chemiluminescent reagent were captured in ChemiDoc MP System and quantified in a linear range using the ImageLab 5.2.1 software (Bio-Rad).

### Immunohistochemistry

Mice were perfused transcardially with PBS and fixed in 4% phosphate-buffered paraformaldehyde before paraffin embedding. Sagittal brain sections (5 μm) were deparaffinized in xylene, rehydrated and microwave heated in citrate buffer (10 mM, pH 6.0). Sections were incubated overnight at 4°C with mouse anti-phosphorylated tau antibody (CP13, 1:50). Sections were then incubated with a biotin-conjugated anti-mouse secondary antibody (1:200) and revealed with the DAB peroxidase substrate kit (Vector laboratories) before imaging with a Nikon Eclipse 80i microscope. For immunofluorescence staining of human hippocampus, frozen human samples (10 µm) were fixed in 4% PFA and heated in citrate buffer. Sections were incubated overnight at 4°C with mouse anti-phosphorylated tau (CP13, 1:50), mouse anti-p62 (1:200) or rabbit anti-LC3 (1:200) antibodies followed by anti-mouse AlexaFluor488 (1:300), biotin-conjugated anti-mouse or rabbit antibodies (1:300), Streptavidin Cy3 (1:1000) and Hoechst (1:10,000). Confocal images (63x; zoom 1) were obtained with a Zeiss Axio Examiner LSM700 laser scanning microscope. All images were analyzed with ImageJ software (NIH). *Gallyas staining* We used the simplified modified Gallyas staining method to stain NFTs (Kuninaka et al., 2015). Brain sections were deparaffinized and rehydrated, submerged in 0,3% KMnO_4_ (10 min), treated with 1% oxalic acid (1 min) and washed with H_2_O_d_. After alkaline silver iodide treatment (1 min), sections were washed (x 3) with 0.5% acetic acid and developed (6-7 min) with solution A (0.2% silver nitrate, 0,2% ammonium nitrate, 1% g tungstosilicic acid and 0.2% formaldehyde) and solution B (5% anhydrous sodium carbonate). Sections were washed (x3) in 1% acetic acid and stained with 0.5% gold chloride. Nuclear fast red (0,1%) was used as counterstaining, and samples were dehydrated and mounted.

### Statistical analysis

Statistical analysis with Prism software (GraphPad, La Jolla, CA) was performed using one- or two-way ANOVA followed by Dunnett’s or Tukey’s *post hoc* test as indicated in the figure legend. For data not normally distributed, Kruskal-Wallis followed by Dunn’s *post hoc* test was used. *P* values less than 0.05 were considered significant. The significance level is indicated using asterisks: * *P* < 0.05, ** *P* < 0.01 and *** *P* < 0.001

## Results

### Pathological tau is accumulated in autophagic compartments in the hippocampus in tauopathies, including familial Alzheimer’s disease

To investigate the link between tau accumulation and autophagy dysfunction in tauopathies, we analyzed the hippocampus of 28 individuals clinically diagnosed with *PSEN*-linked familial AD (FAD; n = 7, mean age: 53.1 ± 5.7), CBD (n = 8, mean age: 70.5 ± 6.7), PiD (n = 6, mean age: 68.3 ± 13.7) and age and sex-matched controls (n = 7, mean age: 51.4 ± 7.6) (**Table 1**). Biochemical analysis showed differential levels of total and phosphorylated (Thr 181, Ser 202, Thr 217, Ser 396/404) tau species (**Fig. 1A,B**). Tau antibody revealed protein bands of ∼60-64, 68 and 72 kDa in FAD samples, whereas 64-68 and 60-64 kDa tau bands were predominant in CBD and PiD, respectively (**Fig. 1A**). This differential biochemical profile of tau bands among tauopathies is due to the accumulation of 3R/4R-tau in FAD, 4R-tau in CBD and 3R tau in PiD (Greenberg and Davies, 1990; Lee et al., 1991; Buee Scherrer et al., 1996; Delacourte et al., 1996). Interestingly, total and phosphorylated tau were elevated in patients harboring FAD-linked *PSEN* mutations compared to other tauopathy patients (FAD *vs* control, Tau: *P* < 0.01; Ser 396/404: *P* < 0.001; Thr 217, *P* < 0.01; Ser 202: *P* < 0.01; Thr 181, *P* < 0.01; **Fig. 1A**), When normalizing phosphorylated tau to total tau levels, and due to elevated total tau levels in all tauopathies, only differences in Ser 396/404 (*P* < 0.001) and Ser 202 (*P* < 0.05) were found (data not shown). To examine whether accumulated tau is associated with autophagic flux failure, we analyzed the lysosomal-associated membrane protein 2A (LAMP2A), the membrane-associated autophagosome protein light chain 3 (LC3) and sequestosome 1 (SQSTM1/p62), a protein that acts as an adaptor for selective autophagic degradation of ubiquitinated cargos. Number of autophagosomes was assessed by measuring LC3-I conversion to LC3-II, whereas LC3-II and p62 levels were quantified to evaluate autolysosomes degradative efficiency (Mizushima et al., 2010). p62 was significantly increased in FAD (*P* < 0.05), CBD (*P* < 0.05) and PiD (*P* < 0.001), and LAMP2A was slightly enhanced in FAD samples (**Fig. 1C**), suggesting disrupted macroautophagy and impaired autophagosome/autolysosome clearance in tauopathies. Levels of LC3-I and LC3-II, and LC3-II/LC3-I ratio were unchanged in tauopathies, but LC3 precursor (proLC3) was significantly increased in FAD (*P*< 0.05), indicating the possibility of altered autophagy flux initiation in these patients (**Fig. 1C**). In agreement, immunohistochemical analyses revealed significant increase of pTau (Ser 202; *P* < 0.05) and p62 (*P* < 0.05), and high colocalization of pTau/p62 and pTau/LC3 in neuropil threads and neuronal soma, respectively, in the hippocampus of FAD, CBD and PiD patients **(Fig. 2A,B; Suppl Fig 1A,B**). Indeed, colocalization analysis revealed a positive correlation between pTau and p62 in the hippocampus of all tauopathy patients (*P* < 0.001; **Fig. 2B**). Biochemical analysis of purified autophagic vesicles from control and FAD hippocampus revealed a significant accumulation of tau and pTau species in autophagosomes, autolysosomes and/or lysosomes in FAD patients compared with controls (*P* < 0.05; **Fig. 2C, Suppl Fig. 1C**). Together, these results suggest that autophagy is compromised in tauopathies leading to pathological tau accumulation in autophagic compartments.

**Figure 1.**
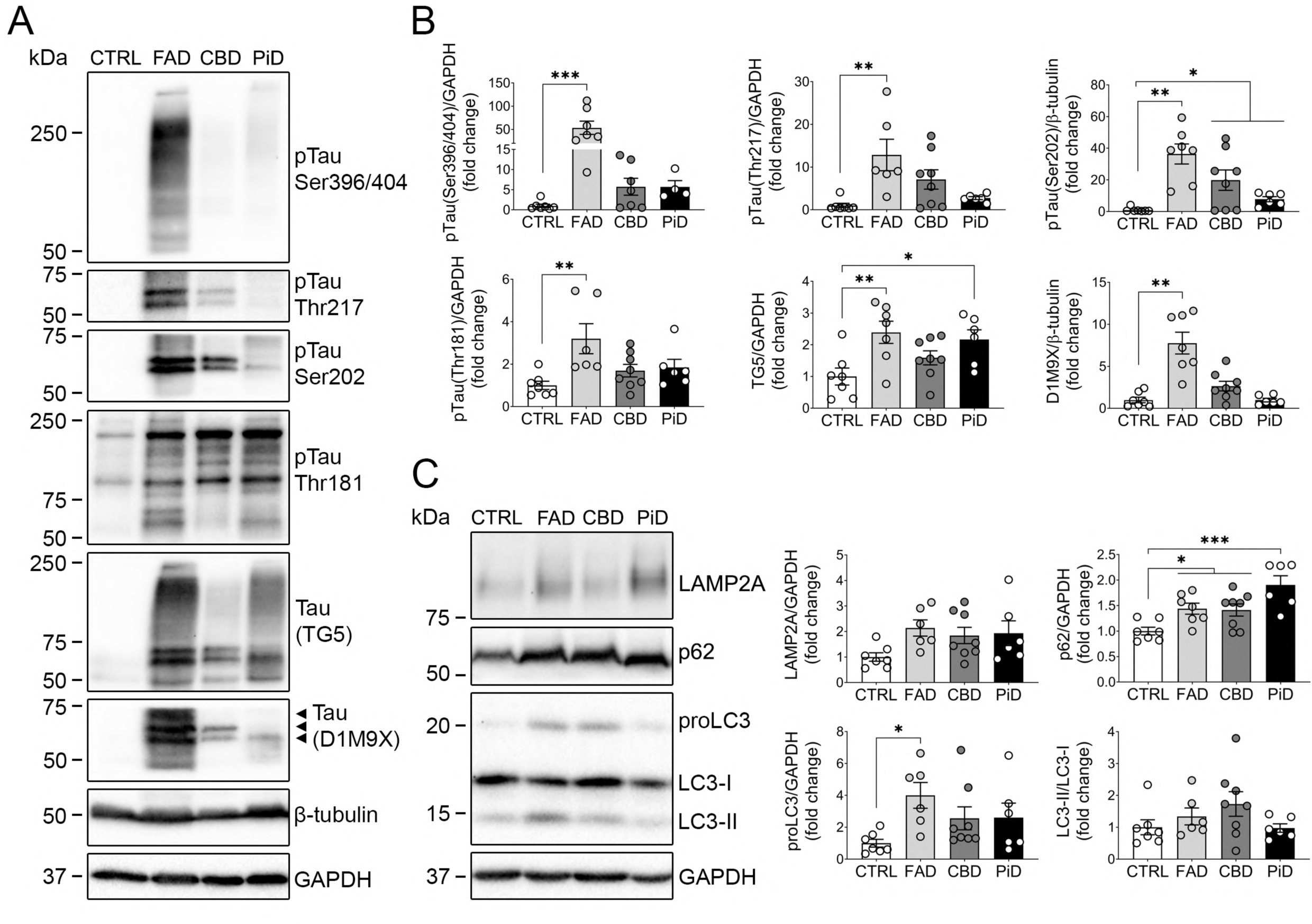
Increased p62 levels are associated with accumulation of total and phosphorylated tau in the hippocampus of tauopathy patients. Western blot and quantification analysis of phosphorylated (p) Ser 393/404 (PHF-1), Thr 217, Ser 202 (CP13) and Thr 181 (AT270), and total tau (TG5 and D1M9X) **(A,B)**, and autophagic/lysosomal markers LAMP2A, p62 and LC3 **(C)** in hippocampal lysates from controls (CTRL) and patients with FAD, CBD and PiD. Arrowheads indicate ∼64, 68 and 72 kDa tau bands. Protein levels were normalized to β-tubulin, GAPDH or total tau. Data represent mean ± SEM of multiple individuals (n = 6-8) per group. One-way ANOVA followed by Dunnett’s post hoc test or Kruskal-Wallis followed by Dunn’s post-hoc test was used as statistical test. \**P* < 0.05, \*\**P* < 0.01, \*\*\**P* < 0.001. CTRL: control; FAD: familial Alzheimer’s disease; CBD: corticobasal degeneration; PiD: Pick’s disease

**Figure 2.**
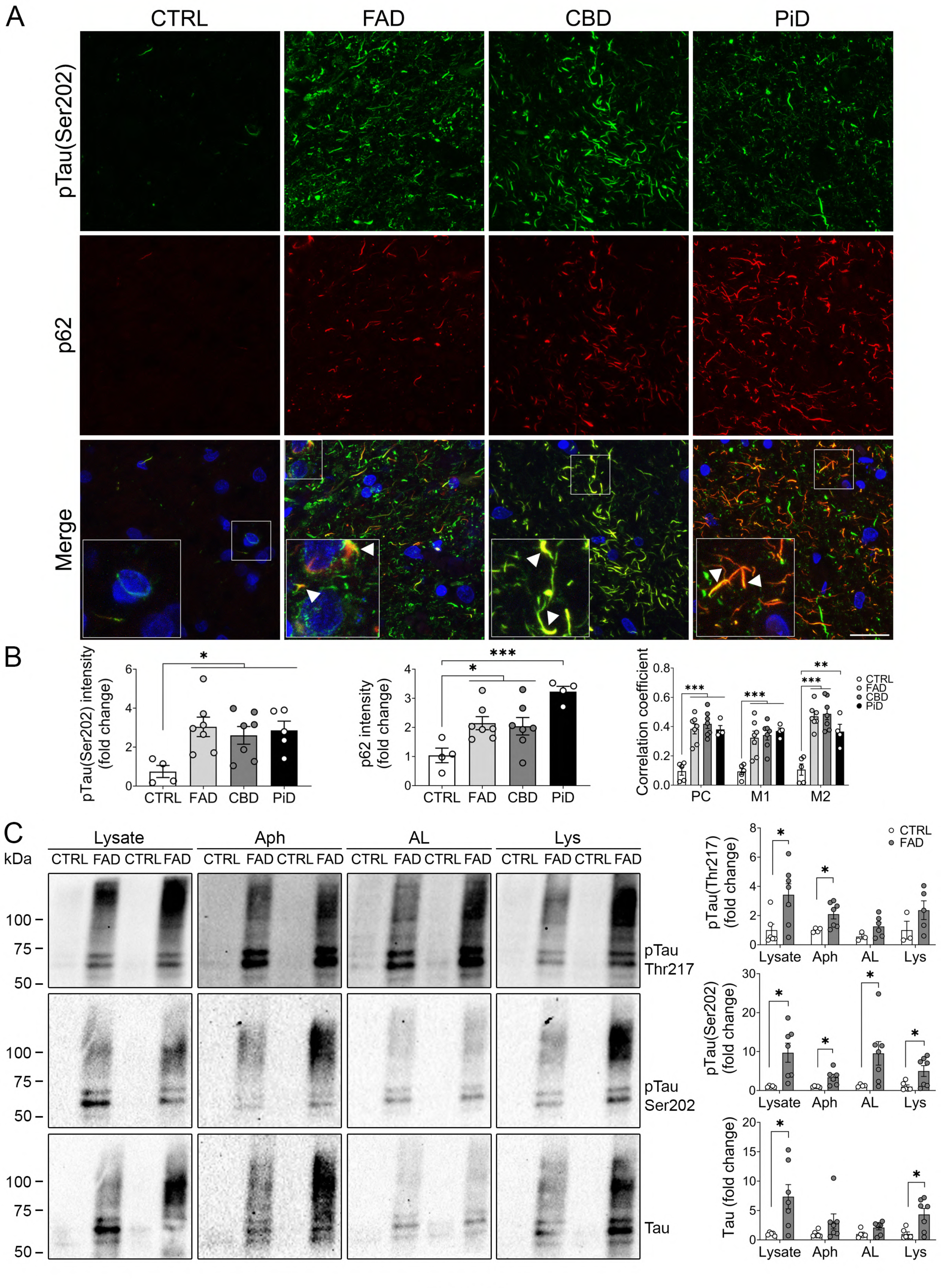
Phosphorylated tau accumulates in autophagic vesicles of hippocampus of tauopathy patients. **A,B**, Representative immunofluorescence images of pTau Ser 202 (CP13, green) and p62 (red) **(A)**, and quantitative analyses of total intensity per area and colocalization of pTau Ser202 and p62 **(B)** in hippocampal brain sections from CTRL and FAD, CBD and PiD patients. Data represent mean ± SEM of multiple individuals (n = 4-7) per group. Scale bar: 20 µm. One-way ANOVA followed by Dunnett’s post hoc test or Kruskal-Wallis followed by Dunn’s post hoc test was used as statistical test. **C,** Western blot images and quantification analyses of pTau (Thr 217 and Ser 202) and total tau (D1M9X) in lysates and purified autophagosomes (Aph), autolysosomes (AL) and/or lysosomes (Lys) from hippocampus of human CTRL and FAD patients. Welch’s t test or Mann-Whitney test were used as statistical test. \**P* < 0.05, \*\**P* < 0.01, \*\*\**P* < 0.001. CTRL: control; FAD: familial Alzheimer’s disease; CBD: corticobasal degeneration; PiD: Pick’s disease; PC: Pearson’s correlation coefficient; M1/M2: Mander’s overlap coefficient 1 and 2.

### FAD-linked PSEN1 mutations enhance pathological tau and LC3-I in human iPSCs-derived neurons

The above results promoted us to examine levels of tau and autophagy markers in human iPSCs-derived hippocampal neurons (60 DIV) from a healthy control donor and a patient harboring the heterozygous *PSEN1* G206D mutation (*PSEN1* exon 7) that causes early-onset FAD (Diaz-Guerra et al., 2019; Rodriguez-Traver et al., 2020). This analysis was performed in the absence or presence of the autophagosome-lysosome fusion inhibitor chloroquine (CQ) (Mauthe et al., 2018). In basal (vehicle) conditions, no significant changes in autophagy markers were observed in *PSEN1* G206D human iPSCs-derived neurons (**Fig. 3A**). As expected, CQ (25 µM) induced an overall increase of p62 (∼1.4 fold), LC3-II (∼2.5-3 fold, data not shown) and LC3-II/LC3-I ratio (∼3 fold) indicating blockade of autophagosome clearance (Treatment effect: *P* < 0.001; **Fig. 3A**). However, p62 was not significantly accumulated in *PSEN1* G206D iPSCs-derived neurons treated with CQ (**Fig. 3A**). In control neurons, CQ treatment significantly reduced LC3-I levels likely due to its conversion to LC3-II (*P* < 0.05), an effect that was not observed in *PSEN1* G206D iPSCs-derived neurons (*P* > 0.05, **Fig. 3A**). This result suggests that patient-derived neurons are not as sensitive to CQ as controls. This effect, along with unchanged LC3-II levels between control and patient neurons in basal and CQ-treated conditions (*P* > 0.05, data not shown), indicates enhanced autophagy flux initiation in *PSEN1* G206D iPSCs-derived neurons. As shown in **Fig. 3B**, we found bands of ∼64-68 kDa monomeric tau; and ∼120-150 kDa bands of high molecular weight tau (HMW-Tau) likely corresponding to tau aggregates/oligomers (Lasagna-Reeves et al., 2010; Patterson et al., 2011). Surprisingly, monomeric tau (∼64-68 kDa) was decreased (*P* < 0.05) and HMW-Tau/Tau was elevated in both basal and CQ conditions in *PSEN1* G206D iPSC-derived neurons (*P* < 0.05 and *P* < 0.01, **Fig. 3B**). CQ-treated control and G206D iPSCs-derived neurons reduced monomeric pTau (Ser 396/404) but enhanced phosphorylated HMW-Tau/pTau aggregates (**Fig. 3B**). CQ treatment enhanced mature glycosylated (mAPP) and immature non-glycosylated (iAPP) APP and APP-αCTFs (*P* < 0.001) in control but not *PSEN1* G206D iPSCs-derived neurons (**Fig. 3B**), suggesting that this mutation reduces the non-amyloidogenic pathway when autophagy is blocked (Jacobsen and Iverfeldt, 2011). In agreement, *PSEN1* G206D iPSCs-derived neurons secrete higher Aβ40 and Aβ42 levels and release lower tau independently of treatments (**Fig. 3C)**. These results suggest that the *PSEN1* G206D mutant affects negatively autophagosome flux and potentiates the amyloidogenic and tau pathways likely increasing intracellular oligomeric tau species in human neurons.

**Figure 3.**
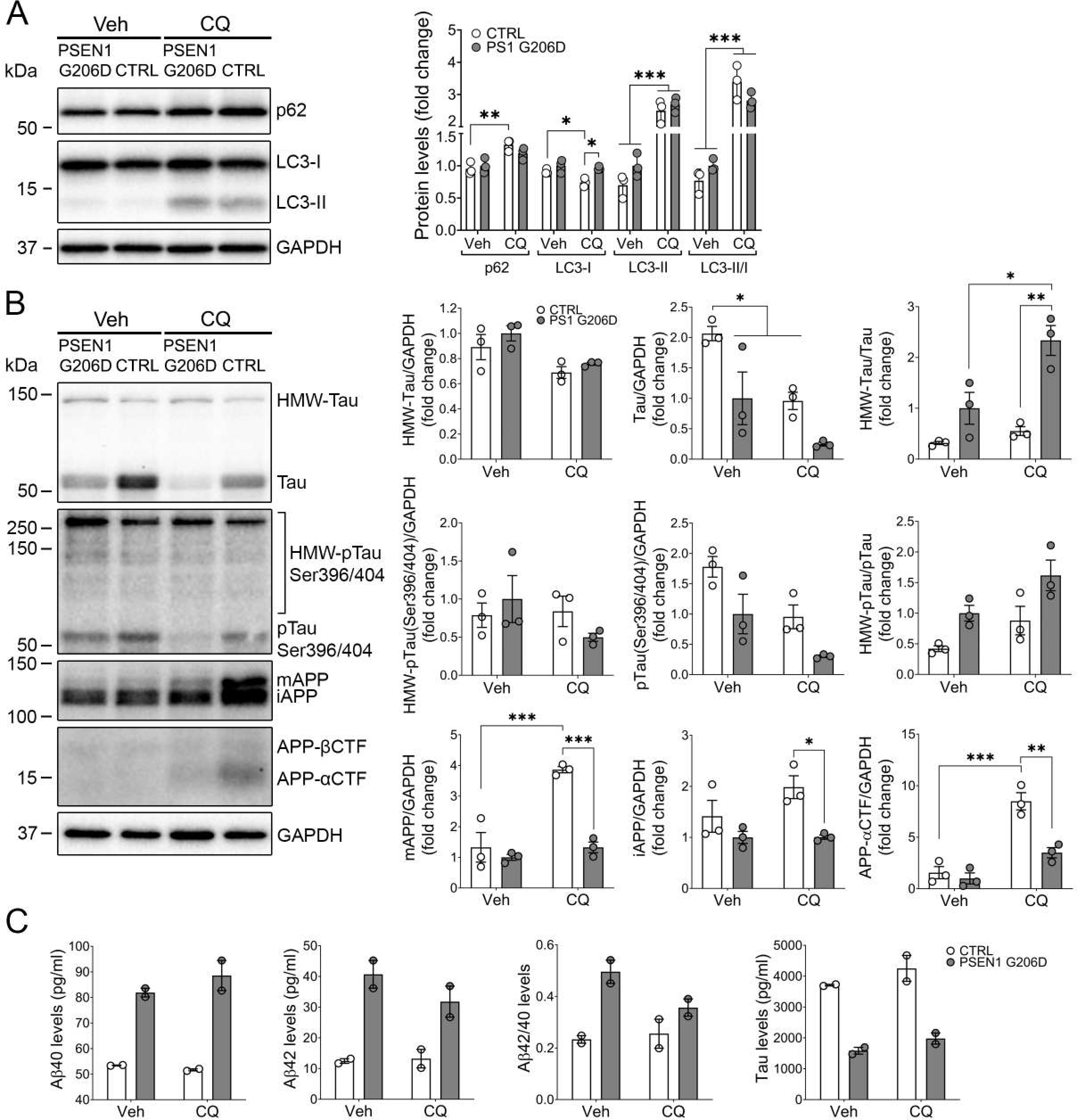
Increased intracellular aggregated Tau and decreased Tau release in human iPSCs-derived neurons expressing the *PSEN1* G206D mutation. **A**, Western blot images and quantification of p62, LC3-I and LC3-II, normalized to GAPDH, in lysates of iPSC-derived neurons treated with vehicle or CQ for 24 h. **B,** Western blot images and quantitative analysis of total tau (D1M9X), phosphorylated (p) Ser 396/404 tau (PHF-1), and APP full length and CTFs in lysates of iPSCs-derived neurons treated with vehicle or CQ for 24 h. **C,** Milliplex measurement of β40 and Aβ42 peptides, and total tau in the cultured media of *PSEN1* G206D iPSCs-derived cultured neurons. Protein levels were normalized to GAPDH, LC3-I, or ∼64-68 kDa tau, as indicated. Data represent mean ± SEM of three biological replicates per condition. Two-way ANOVA followed by Tukey’s post hoc test was used as a statistical test. \**P* < 0.05, \*\**P* < 0.01. \*\*\**P* < 0.001. HMW: high molecular weight; mAPP: mature (glycosylated) APP; iAPP: immature (non-glycosylated) APP.

### FAD-linked PSEN1 mutations block autophagic flux in human primary fibroblasts

To examine whether the effect of FAD-linked *PSEN1* mutations in autophagy is tau-dependent, we monitored autophagy dynamics in primary human skin fibroblasts that lack tau expression derived from healthy controls (CTRL 1 and 2) and patients harboring *PSEN1* G206D and L286P (*PSEN1* exon 8) missense mutations (Diaz-Guerra et al., 2019). Treatment of fibroblasts with CQ (10 μM) for 24 h increased autophagic vacuoles, visualized under a bright-field microscope (not shown), and elevated p62 (*P* < 0.001), LC3-II (*P* < 0.001) and LC3-II/LC3-I ratio (*P* < 0.001; **Fig. 4A**). Fibroblasts expressing the G206D and L286P *PSEN1* mutations showed similar p62 levels in basal or CQ conditions compared to controls (*P* > 0.05, **Fig. 4A**). Interestingly, LC3-I and -II were significantly elevated in human fibroblasts expressing FAD-linked *PSEN1* mutants in basal conditions (G206D, *P* < 0.01; L286P, *P* < 0.05), an effect not observed after CQ treatment (**Fig. 4A**). The increase of LC3 suggests potentiation of autophagosomes formation and/or altered autolysosomes clearance in FAD-linked *PSEN1* fibroblasts. We next evaluated autophagy flux in primary fibroblasts transiently expressing the mCherry-EGFP-LC3 reporter, which allows monitoring of biogenesis of autophagosomes (mCherry+/EGFP+, yellow) and autolysosomes (mCherry+/EGFP-, red) due to quenching of EGFP fluorescence (Mizushima et al., 2010) (**Fig. 4B)**. In basal conditions, autophagosome number (yellow puncta) were unchanged and autolysosomes (red puncta) were increased in fibroblasts expressing the *PSEN1* G206D (54.9 ± 7.0; *P* < 0.05) and L286P (68.1 ± 12.3; *P* < 0.001) mutants **(Fig. 4C,D**). CQ induced an increase of autophagosomes and autolysosomes in control fibroblasts, whereas their number were increased or maintained, respectively, in *PSEN1* G206D and L286P fibroblasts. The observed increase of total LC3 and autolysosomes indicate disrupted autophagy flux probably due to decreased lysosomal activity in *PSEN1* G206D and L286P mutant fibroblasts.

**Figure 4.**
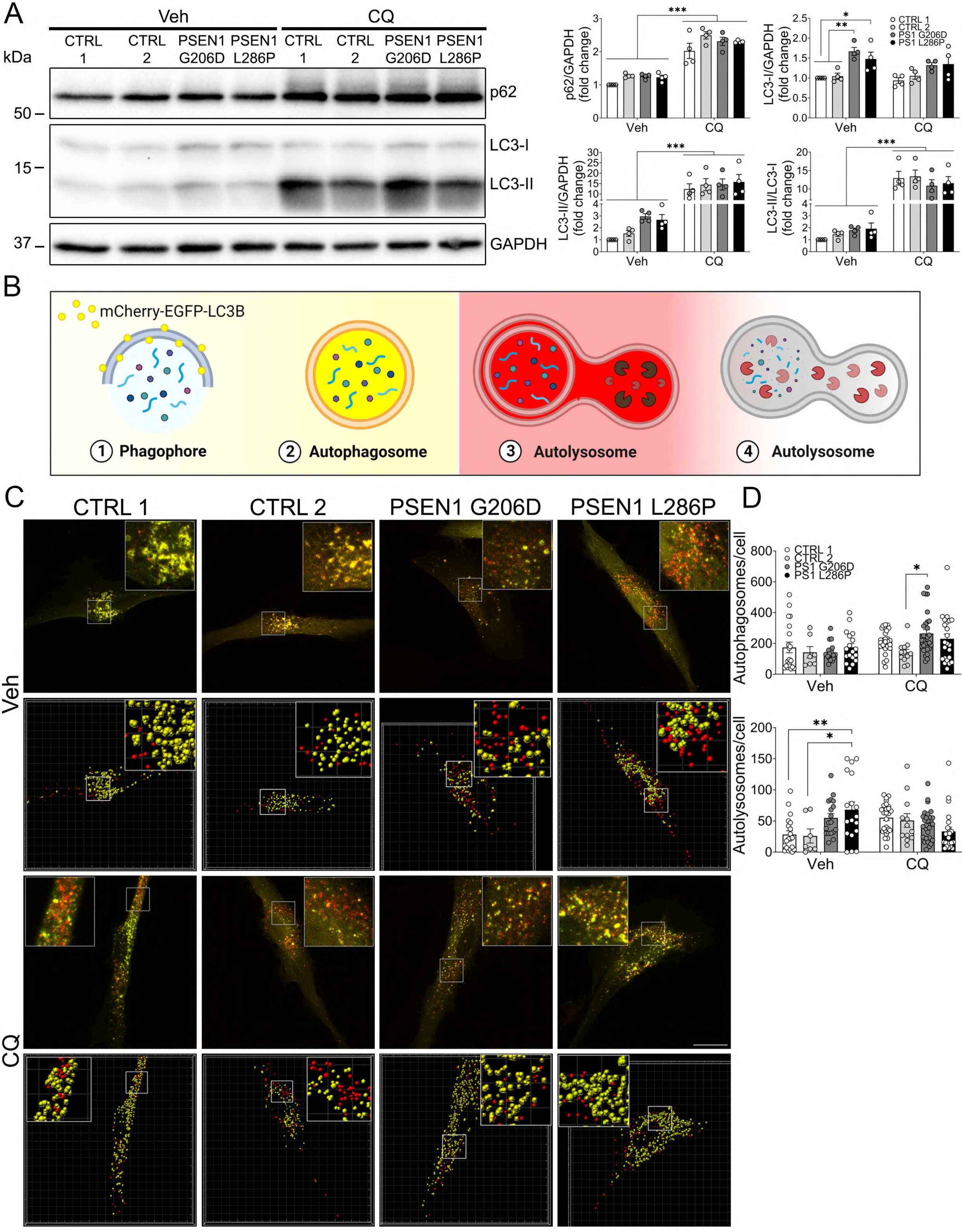
FAD-linked *PSEN1* G206D and L286P mutations impair autolysosomes clearance in primary human fibroblasts. **A**, Western blotting and quantitative analysis of p62, LC3-I and LC3-II protein levels in lysates of cultured human fibroblasts treated with vehicle or CQ for 24 h. Protein levels were normalized to GAPDH or LC3-I, as indicated. Data are mean ± SEM of independent cultures (n = 4). Two-way ANOVA followed by Tukey’s post hoc test was used as a statistical test. \**P* < 0.05, \*\**P* < 0.01, \*\*\**P* < 0.001. **B,** Scheme of the autophagy tandem sensor mCherry-EGFP-LC3B. Autophagosomes are labelled in yellow due to mCherry/EGFP co-staining, and after fusing with lysosomes EGFP is quenched by the acidic pH and autolysosomes are labelled in red (mCherry+). **C, D,** Representative confocal microscopy (first and third rows) and Imaris software (second and fourth rows) images (**C**) and quantitative analysis (**D**) of autophagosomes (yellow puncta) and autolysosomes (red puncta) per cell in primary human fibroblasts expressing mCherry-EGFP-LC3B and treated with vehicle (top) or CQ (bottom) for 24 h. Scale bar: 20 μm. Data are mean ± SEM of multiple cells per condition (n = 7-29) from 6 independent cultures. Two-way ANOVA followed by Tukey’s post hoc test was used as a statistical test. * *P* < 0.05, ** *P* < 0.01.

### Loss of neuronal PS enhances phosphorylated and aggregated tau in the hippocampus of Tau transgenic mice

To investigate further the role of PS/γ-secretase on tau pathology *in vivo*, we next analyzed novel PS1 cKO;Tau (*PS1* f/f;CamKIIα-Cre;Tau) and PS cKO;Tau (*PS1* f/f;PS2^-/-^;CamKIIα-Cre;Tau) mice expressing the FTD-linked P301S tau and lacking *PS1* or both *PS* in excitatory neurons of the postnatal forebrain (Soto-Faguas et al., 2021). Biochemical analyses show similar total human tau levels (∼4-fold change) in all Tau transgenic groups, whereas pTau at Thr 217, Ser 202 and Thr 181 were enhanced (∼3-20-fold) in the hippocampus of 6 month-old PS1 cKO;Tau and/or PS cKO;Tau mice compared to WT mice (one-way ANOVA, Thr 217: *P* < 0.001; Ser 202: *P* < 0.01; Thr 181: *P* < 0.01; **Fig. 5A**). pTau species, particularly at Ser 202 and Thr 217, are slightly increased in PS cKO;Tau mice compared to Tau and/or PS1 cKO;Tau mice (*P* < 0.05; **Fig. 5A**). Immunostaining of pTau (Ser 202) confirmed increased somatic and neuronal projections staining and pathological tau-positive cells in the hippocampus (one-way ANOVA, genotype effect: CA3, *P* < 0.05; DG, *P* < 0.01;), retrosplenial cortex (*P* < 0.01) and amygdala (*P* < 0.01) of PS1 cKO;Tau and PS cKO;Tau mice compared to control and/or Tau mice (**Suppl Fig. 2**). Interestingly, Gallyas staining revealed significant increase of conformational aggregated tau in hippocampus and EC of PS cKO;Tau mice at 6 months compared with control and Tau mice (CA3: *P* < 0.05; EC: *P* < 0.01; **Fig. 5B)**. These results indicate that loss of *PS* function in neurons increases human tau phosphorylation and aggregation.

**Figure 5.**
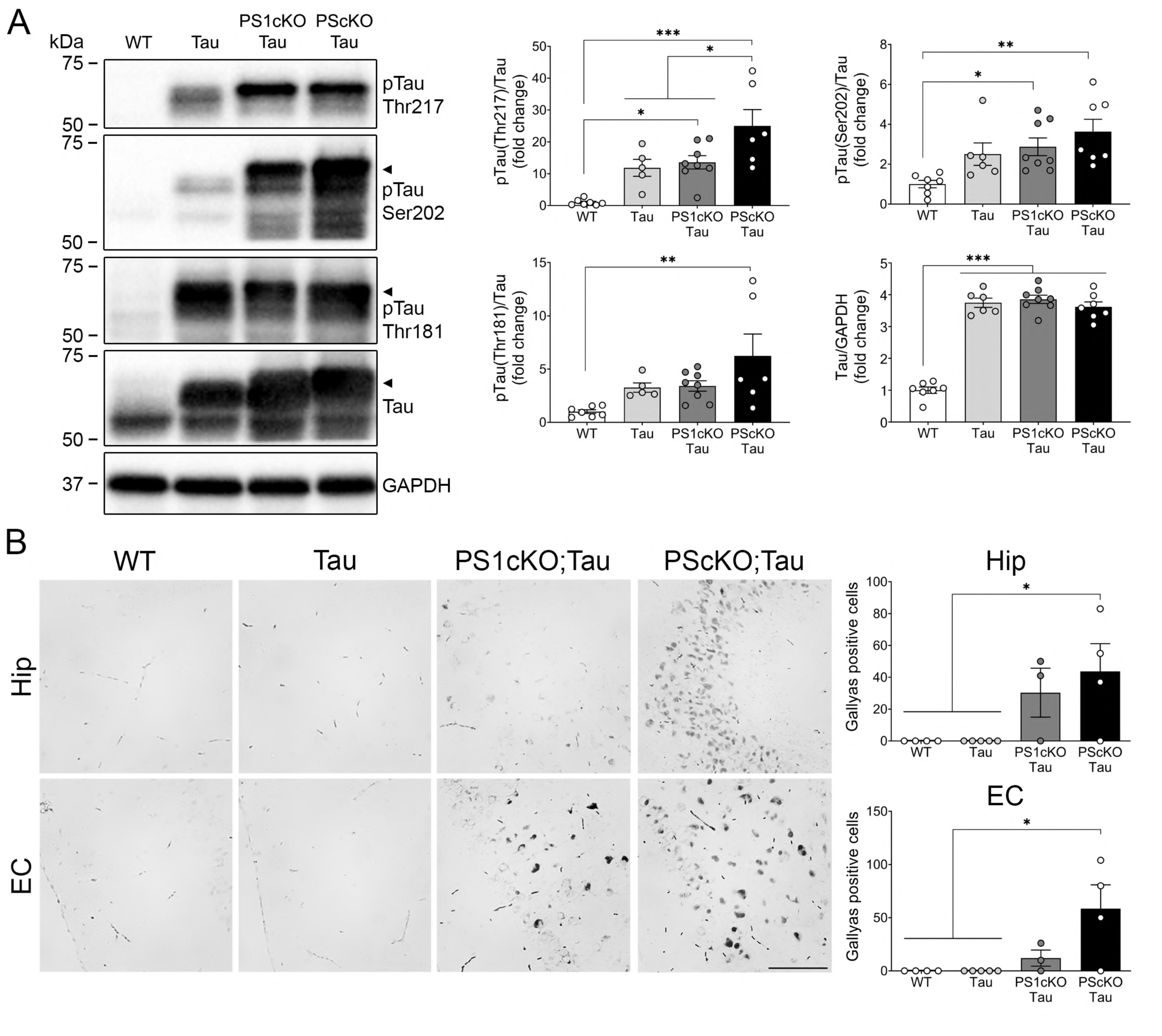
Loss of neuronal PS enhances cerebral phosphorylated and aggregated human tau. **A**, Western blot analysis of phosphorylated (p) Ser 202 (CP13), Thr 217, Thr 181 (AT270) and total tau (D1M9X) in the hippocampus of 6 month-old wild-type (WT) and tau transgenic mice (Tau) lacking neuronal PS1 (PS1 cKO;Tau) or both PS (PS cKO;Tau). Protein levels were normalized to GAPDH or total tau, as indicated. Values represent mean ± SEM (n=7-8 mice/group). **B,** Representative images and quantification of Gallyas-stained neurons in coronal sections of CA3 hippocampus (Hip) and entorhinal cortex (EC) of 6 month-old control (WT), Tau, PS1cKO;Tau and PScKO;Tau transgenic mice. Scale bar: 100 µm. Values represent mean of Gallyas+ cells/section ± SEM (n=3 slices per mice, 4 mice/group). One-way ANOVA followed by Tukey’s post hoc test was used as statistical test. \**P* < 0.05, \*\**P* < 0.01, \*\*\**P* < 0.001.

### PS is required for autophagy-mediated tau degradation in neurons

Considering that pathological tau is eliminated via autophagy (Wang et al., 2009; Kruger et al., 2012; Silva et al., 2020), and PS regulates the autophagic-lysosomal system (see above, (Lee et al., 2010; Neely et al., 2011; Lee et al., 2015)), it is plausible that PS-dependent tau homeostasis could involve autophagy deregulation. Biochemical analyses of autophagy-related and upstream molecules in hippocampus of 6 month-old Tau, PS1cKO;Tau and PScKO;Tau mice revealed significant increases of p62 (*P* < 0.001), active pAkt (Thr308), Akt-mediated phosphorylated GSK3β (Ser 9; *P <* 0.05) and proline-rich Akt substrate (PRAS40, Thr 246; *P* < 0.001), and total and phosphorylated (Ser235/236) S6, but not changes in LC3-II/I ratio and pmTOR (Ser 2448), or its downstream phosphorylated effectors ULK1 (Ser 757) and p70S6K (Thr 389), pAMPK (Thr 172) and pAkt (Ser 473), in PScKO;Tau mice (**Figs. 6A,B**). Additionally, significant increases of LC3-II/I ratio, pAkt, pPRAS40, and total and phosphorylated S6, and decreased p-p70S6K (Thr 389), were detected in the hippocampus of 6 month-old PScKO mice (*P* < 0.01, **Fig. 6C**). Hippocampal levels of pULK1 (Ser757) and pAkt (Thr 308) were increased, but not significantly, in PS cKO mice compared with control group. These results suggest increased mTORC1 activity and, consequently, inhibition of autophagy initiation in non- and tau-transgenic PS cKO mice. In agreement with autophagy-related proteins changes in PScKO mice, we found significant accumulation of endogenous pTau (Ser 202) in total lysates and purified autophagosomes and lysosomes, and reduced total tau in autolysosomes, of PScKO mouse hippocampus (*P* < 0.05; **Fig. 6D**). Together, these findings indicate that neuronal PS is required for efficient autophagy-mediated tau degradation in neurons through a dual mechanism involving autophagy induction via blockage of Akt/PRAS40-dependent mTORC1 activation and promoting autophagosome/lysosome fusion.

**Figure 6.**
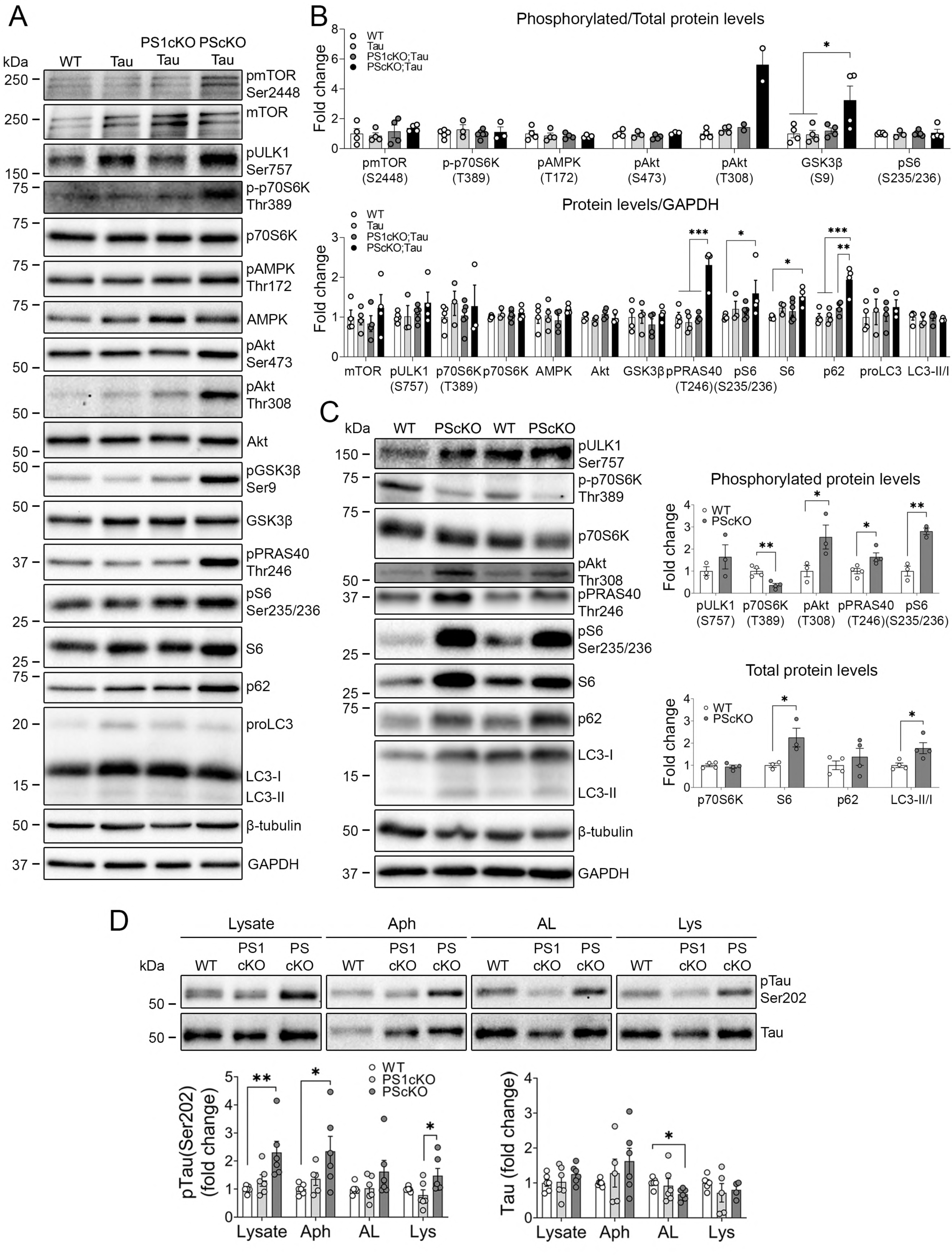
Loss of neuronal PS inhibits autophagy induction by increasing mTOR activity. **A, B** Biochemical analysis and quantification of phosphorylated and/or total mTOR, ULK1, p70S6K, AMPK, Akt, GSK3β, PRAS40, S6, p62 and LC3 in hippocampal lysates of 6 month-old WT, Tau, PS1 cKO;Tau, PS cKO;Tau transgenic mice. Phosphorylated proteins were normalized to their corresponding total proteins (except for pULK1 and pPRAS40), and total proteins were normalized to GAPDH, except for pPRAS40 that was normalized to β-tubulin. **C**, Similar biochemical analysis of autophagy modulators in hippocampal lysates of 6 month-old control (WT) and PS cKO mice. Protein levels were normalized to GAPDH or β-tubulin except for p-p70S6K (Thr389) that was normalized to total p70S6K. **D**, Analysis of endogenous tau and pTau (Ser 202, Thr217) in purified hippocampal autophagosomes, autolysosomes and lysosomes of control (WT), PS1cKO and PScKO mice. Values represent mean ± SEM (n=3-5 mice/group). Statistical analysis was determined by one-way ANOVA followed by Tukey’s post hoc tests. \**P* < 0.05, \*\**P* < 0.01, \*\*\**P* < 0.001

### Proteasome-dependent regulation of neuronal accumulation and release of tau requires functional PS

p62 acts as a selective autophagy adaptor protein of ubiquitinated cargos. To further investigate the link between PS-dependent tau pathology and proteasome/autophagy pathways, we established a novel *in vitro* tauopathy model of primary cortical neurons from tau transgenic (Tau+) embryos harboring floxed *PS1*;*PS2*^+/+^ or *PS1*;*PS2*^-/-^ alleles to silence conditionally PS1 or both PS using active or mutant Cre-recombinase lentiviral vectors. PS silencing was biochemically confirmed by reduced PS1 N-terminal fragment (*P* < 0.001, **Fig. 7A**). To study tau levels in response to autophagy and/or proteasome inhibition, primary cortical neurons were treated with CQ or MG132, respectively. Blockade of autophagosome/lysosome fusion efficiently increased p62 and LC3-II/LC3-I ratio compared to vehicle (*P* < 0.001, **Fig. 7A,C**), while proteasome blockage enhanced p62 (*P* < 0.001) without changes in LC3-II/LC3-I (**Fig 7A,C**). Unexpectedly, in contrast to CQ treatment, inhibition of proteasome alone or together with autophagy reduced intracellular phosphorylated and/or total tau levels independently of PS expression (treatment effect: *P* < 0.001, **Fig. 7A,B**). Analysis of culture medium revealed significant elevation of secreted human tau in MG132 and CQ/MG132-treated Tau and PS1cKO;Tau neurons (*P* < 0.001), an effect that was significantly reduced in PScKO;Tau neurons (**Fig. 7D**). Indeed, intraneuronal tau was negatively correlated with extracellular tau in Tau (r = −0.981, *P* < 0.05), PS1cKO;Tau (r = −0.943, *P* < 0.05) and PScKO;Tau (r = −0.956, *P* < 0.05) mice, suggesting that inhibition of proteasome decreases intracellular tau by increasing its release through a mechanism that requires functional PS (**Figs. 7E,F**).

**Figure 7.**
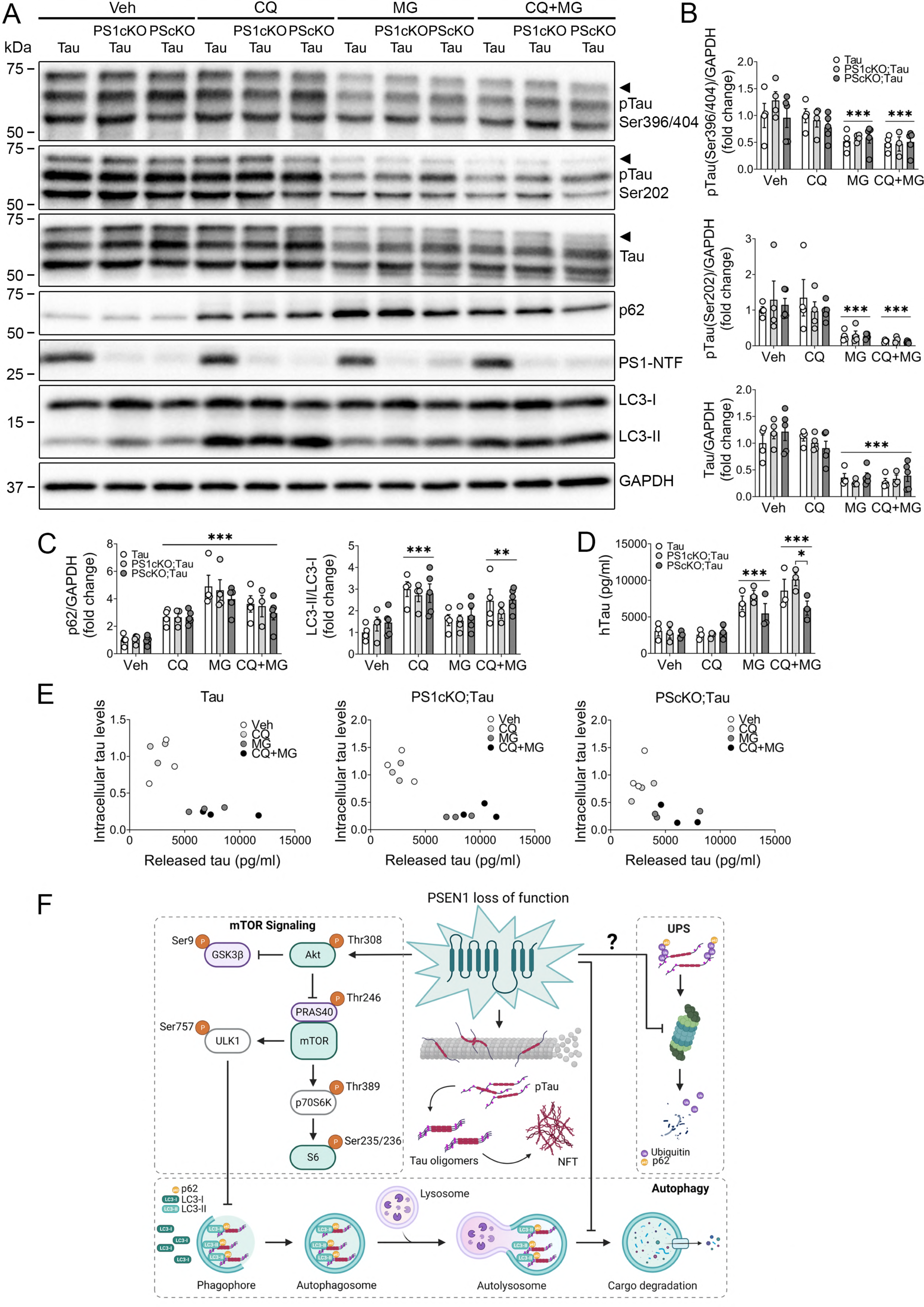
PS mediates tau release after proteasome inhibition in primary cortical neurons. **A-C**, Western blot images and quantitative analyses of phosphorylated (p) tau Ser 396/404 (PHF-1) and Ser 202 (CP13), total tau (D1M9X), PS1-NTF, and autophagy markers p62 and LC3 of lysates of primary cortical neurons (12 DIV) treated with vehicle (Veh), cloroquine (CQ) and/or MG132 (MG) for 24 h. Protein levels were normalized to GAPDH, and values represent mean ± SEM (n = 4-5 embryos/group of 3 independent cultures). **D,** Human (h) tau levels measured by ELISA in the cell culture medium of primary cortical neurons (12 DIV) treated with Veh, CQ and/or MG132 for 24 h. Statistical analysis was determined by two-way ANOVA followed by Tukey’s post hoc tests. \**P* < 0.05, \*\**P* < 0.01, \*\*\**P* < 0.001. **E,** Correlation analyses of intracellular and released hTau levels in primary cortical neurons overexpressing human tau and lacking PS1 (PS1 cKO;Tau) or both PS (PS cKO;Tau), and treated with Veh, CQ and/or MG132 for 24 h. Data was analyzed using two-tailed Pearson’s (r) correlation coefficient. **F**, PS1-dependent molecular pathways that regulate autophagy-mediated tau degradation. PS1 mediates autophagy initiation and autolysosome clearance, whereas loss of PS1 function blocks autophagy induction, by activating mTOR signaling, and autophagy flux resulting in accumulation of pTau. FAD-linked autosomal dominant *PSEN1* mutations increase pathological pTau and aggregated oligomeric tau species in autolysosomes due to defective lysosomal acidification. Besides autophagy, soluble monomeric tau is degraded via the ubiquitin-proteasome system (UPS) reducing its release, a mechanism that requires functional PS.

## Discussion

Inclusions of aggregated hyperphosphorylated tau are pathological hallmarks of sporadic and genetic neurodegenerative tauopathies caused by autosomal dominant mutations in *MAPT* or *PSEN* genes (Raux et al., 2000; Dermaut et al., 2004a; Bernardi et al., 2009). FAD-linked *PSEN1* mutations affect processing of APP and generation of Aβ peptides (Citron et al., 1997; Chavez-Gutierrez et al., 2012; Petit et al., 2022), but **how these pathogenic mutations drive pathological tau accumulation and aggregation remains still unclear**. Understanding the mechanisms linking PS and tau pathology is fundamental for elucidating novel biomarkers and/or targets to develop novel therapeutic interventions in AD. In this study, we show for the first time that PS regulate proteasome/autophagy-mediated tau elimination, and that familial AD-linked *PSEN* mutations may cause progressive tau pathology via a loss of function mechanism on autophagy (**Fig. 7F**).

Our result showing AD-related phosphorylated aggregated tau species in autophagosomes, autolysosomes and/or lysosomes and increased autophagy markers in the hippocampus of tauopathy patients, including those harboring FAD-linked *PSEN1* mutations, suggests disruption of autophagy-mediated tau elimination in tauopathies. Particularly, FAD-linked *PSEN1* G206D and L286P mutations impaired autolysosome clearance in human fibroblasts, probably due to deficient lysosomal acidification, clustering or enzymatic activity as previously reported in fibroblasts (Coffey et al., 2014; Costa-Laparra et al., 2023). Increased proLC3 and LC3-I levels in FAD patients could suggest blockage of autophagy initiation and impaired LC3-I to LC3-II conversion, or enhancement of autophagy initiation to compensate for the defective lysosomal degradative activity. However, this compensatory mechanism does not prevent accumulation of phosphorylated aggregated tau in human iPSCs-derived neurons expressing mutant *PSEN1* G206D. This is in line with recent studies showing that human neurons carrying FAD-linked *PSEN1* mutations show altered autophagy flux and impaired autophagosome clearance leading to increase of Aβ_42_ and pTau (Li et al., 2020; Chong et al., 2022). These findings indicate that **FAD-linked *PSEN1* mutations disrupt autophagy flux by impairing autolysosome clearance**, which likely contributes to intracellular accumulation of aggregated hyperphosphorylated tau (**Fig. 7F**).

### It is interesting that accumulation of pathological phosphorylated tau occurs in autophagic vesicles in hippocampus of *PSEN1*-linked FAD patients and PS-deficient mice

Accumulation of tau in autophagosomes/autolysosomes could result from disruption of autophagy flux as suggested by several studies indicating that loss of PS and pathogenic mutations impair autophagy and lysosome fusion by deregulating autophagy-related genes and/or lysosome acidification (Lee et al., 2010; Neely et al., 2011; Dobrowolski et al., 2012; Reddy et al., 2016; Chong et al., 2018). PS1 deficiency results in reduced autophagosome formation and autophagy-lysosome related genes by decreasing ERK/CREB signaling and increasing GSK3β in human neural stem cells (Chong et al., 2018). Whether PS regulate autophagy flux via γ-secretase-dependent or independent mechanisms is still controversial (Neely et al., 2011; Hung and Livesey, 2018). Based on its role on autophagy-mediated protein degradation, PS1 promotes, independently of γ-secretase activity, autophagic elimination of long-lived proteins, including βCTFs, whereas inhibition of γ-secretase or FAD-linked autosomal mutations disrupt lysosome and autophagosome function via an APP-dependent mechanism (Neely et al., 2011; Bustos et al., 2017; Hung and Livesey, 2018). The effect of PS on mediating autophagy flux by promoting autophagosome/lysosome fusion is further supported by the elevation of LC3-I and/or LC3-II in PS-deficient neurons and mutant *PSEN1*-derived fibroblasts. By contrast, p62 and LC3-II/LC3-I were not affected in the hippocampus or cultured neurons from PScKO;Tau mice suggesting that human tau overexpression may affect autophagy flux, as recently reported in heterologous cells (Feng et al., 2020). Nonetheless, the negative effects of PS deficiency and FAD-linked mutations on autophagy flux are similar indicating that these mutations may act through a loss of function mechanisms in autophagy.

### Besides playing a critical role on autophagy flux by maintaining the acidification of lysosomes and their fusion with autophagosomes, PS may induce autophagy initiation by inhibiting mTOR signaling

PS deficiency resulted in an increase of active pAkt and Akt-mediated PRAS40 phosphorylation, which leads to its dissociation from mTORC1 complex and mTOR activation, as revealed by elevated pS6 and slight increase of its downstream effector pULK1, despite unchanged phosphorylated p70S6K (**Fig. 7F**). This is interesting because it may suggest that mTOR activity depends highly on complex dissociation. Of relevance, increased levels of activated phosphorylated mTOR was previously linked to tau hyperphosphorylation in AD brain (Li et al., 2005; Sun et al., 2014). By contrast, a role of PS1 in restricting autophagy initiation was suggested by reduced mTOR, phosphorylated-mTOR and its complex proteins, and their excessive attachment to lysosomes in PS-deficient fibroblasts (Neely et al., 2011; Reddy et al., 2016). Interestingly, **tau phosphorylation and aggregation is accompanied by p62 accumulation in the hippocampus of FAD-linked *PSEN1* carriers and PS cKO and PS cKO;Tau mice** (**Figs. 1, 5** and (Soto-Faguas et al., 2021)), suggesting disruption of autophagy and/or proteasomal-mediated tau degradation. By delivering ubiquitinated proteins for degradation, p62 is a central modulator of autophagy and ubiquitin-proteasome systems (Nedelsky et al., 2008; Liu et al., 2016), whereas accumulated p62 in the cytosol inhibits the proteasome when autophagy is impaired (Chesser et al., 2013). By contrast, the differential effect of PS in LC3 levels depending of the cellular system (iPSC-derived neurons, fibroblasts and brain) seems to indicate that alternative degradation mechanisms are involved in PS-dependent tau elimination. Nonetheless, monomeric soluble tau is mainly degraded via proteasome whereas oligomeric and aggregated tau is degraded by the autophagy-lysosome system (Lee et al., 2013). Considering that proteasome inhibition mitigates intraneuronal tau and elevates PS-dependent tau secretion, it is plausible that monomeric tau is degraded mainly via the ubiquitin-proteasomal system, and when is blocked tau is released from neurons.

### A direct consequence of disruption of autophagy flux caused by PS dysfunction could be the abnormal secretion of tau

The fact that proteasome inhibitors, either in the presence or absence of functional autophagy, efficiently reduce intracellular tau while increasing its release in a PS-dependent manner, it may have important biological and therapeutic implications. Autophagy inhibitors and activators enhance or mitigate, respectively, pathological tau progression and toxicity in cellular and tau transgenic mouse models (Hamano et al., 2008; Kruger et al., 2012; Schaeffer et al., 2012; Ozcelik et al., 2013; Silva et al., 2020). Indeed, a recent study pointed that p300/CBP-mediated inhibition of autophagic flux promotes tau secretion and propagation in neurons (Chen et al., 2020). Considering that PS tightly regulates CBP (Marambaud et al., 2003; Saura et al., 2004), it is possible that PS could control tau accumulation and/or secretion by affecting CBP-dependent autophagy. Nevertheless, since autophagy induction may also enhance unconventional tau secretion (Mohamed et al., 2014; Kang et al., 2019), the therapeutic potential of autophagy modulators for tauopathies needs to be reconsidered. Considering the impaired autophagosome clearance of tau in FAD-linked *PSEN1* cases, and that the autophagy activator rapamycin increases autophagosomes and lysosomes in PS-deficient fibroblasts (Neely et al., 2011), it is possible that a pharmacological therapeutic approach to enhance lysosomal degradative activity could be more suitable than autophagy induction in FAD.

## Conclusions

Altered proteolytic pathways are associated with accumulation of protein aggregates in age-related neurodegenerative diseases, including FAD. Our results reinforce a key role of PS on autophagy/proteasome-mediated tau degradation in neurons, and its alteration may contribute to tau release and propagation in the degenerative brain. At the mechanistic level, PS dually modulates autophagy-mediated tau degradation in human and mouse neurons by regulating autophagy flux initiation and autolysosome clearance. These autophagy flux alterations along with altered PS-dependent proteasome-mediated tau degradation may contribute to progression of tau pathology in tauopathy dementias. These results may have important implications for translational research on pathological mechanisms underlying tau pathology in vulnerable memory-related neural circuits in AD. In addition, these findings represent a new perspective on the mechanisms linking pathological tau accumulation and autophagy/proteasome deregulation in tauopathies, including FAD, which are important for the development of new therapeutic targets in these devastating disorders.

## Declarations

### Ethics approval

Studies involving human samples were previously approved by the Human Ethical Committee of the Hospital Clinic-IDIBAPS Biobank and Universitat Autònoma de Barcelona.

All animal procedures conducted in the framework of this project were performed according to the European Union regulations (86/609/EEC, 2010/63/EU) in accordance with guidelines and protocols approved by the Animal and Human Ethical Committee (CEEAH) of the Universitat Autònoma de Barcelona and local government of Catalunya (CEEAH/DMAH: 2895/10571 and 4750/10839). Procedures were conducted to reduce the number, pain, suffering, and distress of experimental animals.

### Consent for publication

Not applicable

### Availability of data and materials

Data and materials, including specific experimental protocol information, are available under request.

### Authors’ contributions

AdSB, CMSF and CAS designed and coordinated the study. AdSB and CMSF designed and performed the biochemical, pathological and immunofluorescence analyses in human and mouse brains and cultured neurons. RV and CV generated, characterized and performed pharmacological treatments of human iPSCs-derived neurons. AdSB, CMSF, RV, JRA AD, CV and CAS discussed and interpreted the data. AdSB, CMSF and CAS wrote the initial manuscript with input of all authors. All authors read and approved the final manuscript.

## Acknowledgements

We thank the Hospital Clinic-IDIBAPS Biobank for providing human samples, and Servei d’Estabulari, Histology and Microscope facilities at Institut de Neurociències-UAB, Meritxell Lara for technical microscopy support, Eva Díaz-Guerra for participating in iPSCs differentiation, and José R. Bayascas for technical antibody advice and sharing reagents.

