## Supplemental Figures for "Presenilin-dependent regulation of tau pathology via the autophagy/proteasome pathway"

**Supplementary Figures**

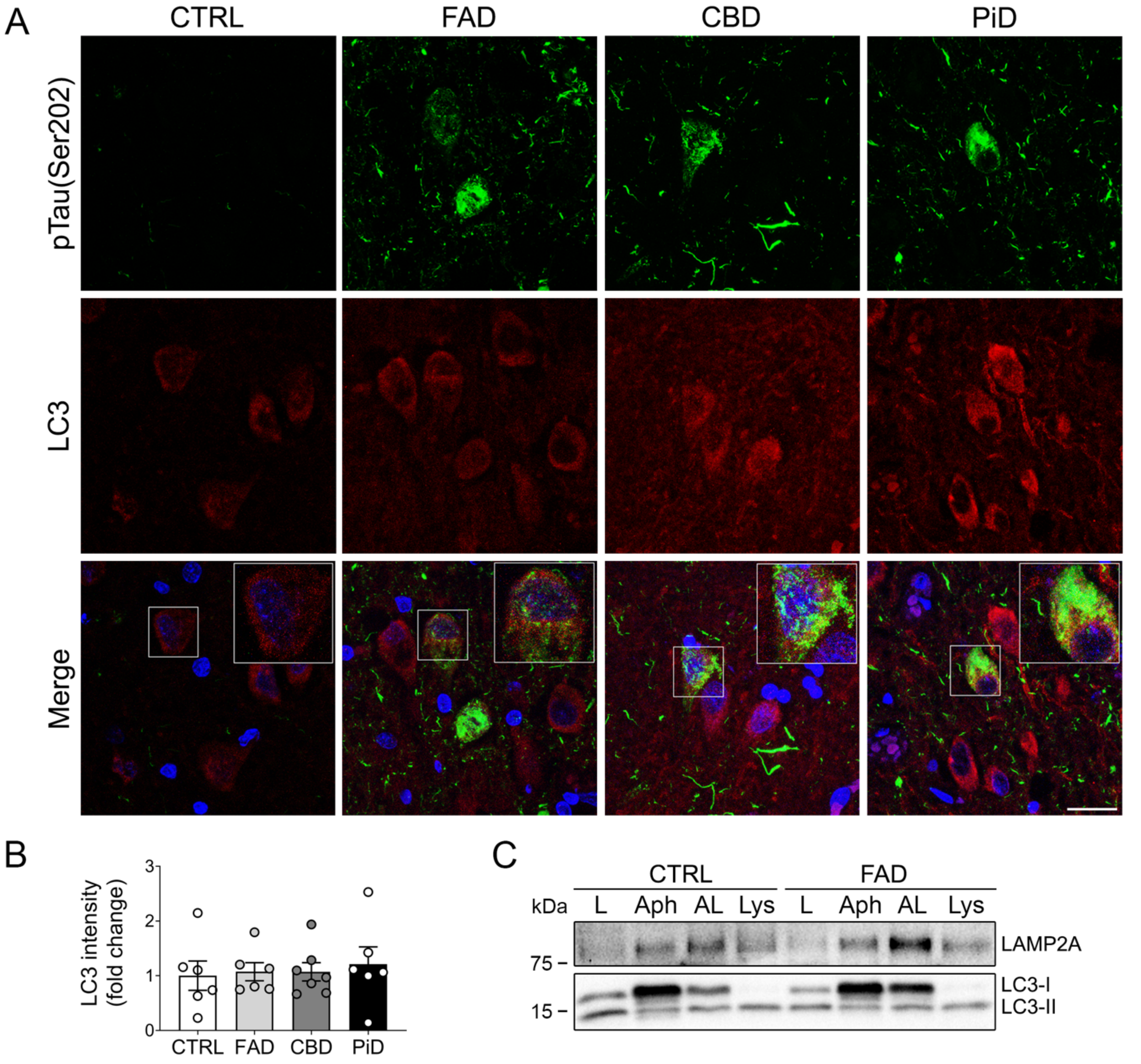


**Supplementary Figure 1. Somatic phosphorylated tau colocalizes with autophagy marker LC3 in the hippocampus of tauopathy patients. A,** Representative immunofluorescence images of hippocampal section from healthy controls (CTRL), FAD, CBD and PiD patients stained with pTau Ser 202 (CP13, green) and LC3 (red). Scale bar: 20 $\mu$m. **B,** Quantitative analysis of total LC3 intensity per area. Data represent mean ± SEM of multiple individuals (n = 6-7) per group. **C,** Biochemical analysis of autophagic and lysosome markers in purified autophagic and lysosomal fractions from control and FAD hippocampus. L: total lysate; Aph: autophagosomes; AL: autolysosomes; Lys: lysosomes.

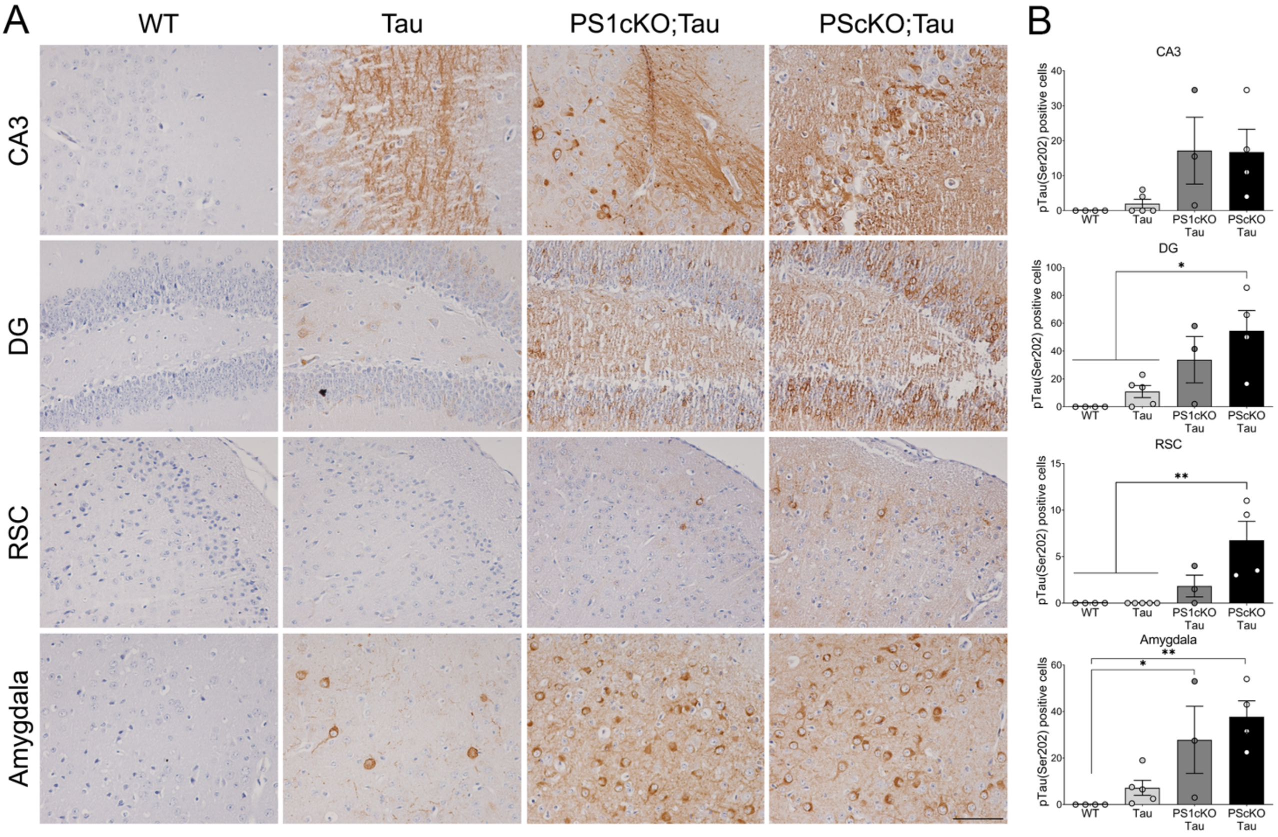


**Supplementary Figure 2. Increased pSer202 tau in brain of PS1 cKO;Tau and PS cKO;Tau mice. A,** Representative immunohistochemical images of phosphorylated tau (pSer202, CP13) in 6 month-old non- (WT) and tau transgenic mice (Tau) lacking neuronal PS1 (PS1 cKO;Tau) or both PS (PS cKO;Tau). Scale bar: 100 $\mu$m. **B,** Quantitative analysis of number of pSer202 positive cells. Values represent mean ± SEM (n=3 slices per mice, 4-5 mice/group). Statistical analysis was determined by one-way ANOVA followed by Bonferroni's post hoc test. *$P < 0.05$, **$P < 0.01$. CA3: CA3 hippocampus; DG: dentate gyrus; RSC: retrosplenial cortex.
